# Zero-Shot, Big-Shot, Active-Shot - How to estimate cell confluence, lazily

**DOI:** 10.1101/2025.01.17.633501

**Authors:** Maximilian Joas, Daniel Freund, Robert Haase, Erhard Rahm, Jan Ewald

## Abstract

Mesenchymal stem cell therapy shows promising results for difficult-to-treat diseases, but standardized manufacturing requires robust quality control through automated cell confluence monitoring. While deep learning can automate confluence estimation, research on cost-effective dataset curation and the role of foundation models in this task remains limited. We systematically investigate the most effective strategies for confluence estimation, focusing on active learning-based dataset curation, goal-specific labeling, and leveraging foundation models for zero-shot inference. Here, we show that zero-shot inference with the Segment Anything Model (SAM) achieves excellent confluence estimation without any task-specific training, outperforming fine-tuned smaller models. Further, our findings demonstrate that active learning does not significantly improve model dataset curation compared to random selection in homogeneous cell datasets. We show that goal-specific, simplified labeling strategies perform comparably to precise annotations while substantially reducing annotation effort. These results challenge common assumptions about dataset curation: neither active learning nor extensive fine-tuning provided significant benefits for our specific use case. Instead, we found that leveraging SAM’s zero-shot capabilities and targeted labeling strategies offers the most cost-effective approach to automated confluence estimation. Our work provides practical guidelines for implementing automated cell monitoring in MSC manufacturing, demonstrating that extensive dataset curation may be unnecessary when foundation models can effectively handle the task out of the box.

## INTRODUCTION

Mesenchymal stem/stromal cells (MSC) are potent Advanced Therapy Medicinal Products (ATMP) and can potentially treat many conditions. MSCs are still not approved for all potential use cases but show promising clinical results in treating degenerative inflammatory diseases, autoimmune diseases, tissue injuries, and chronic degenerative disorders ^1 2^. This is especially relevant because, for diseases like rheumatic arthritis there are no other sufficient treatment options ^3^, and around 18 million people suffer from this disease world-wide ^4^. The starting material for MSCs is sparse and comes from various tissues, like bone marrow, adipose, or umbilical cord tissue. The density of the cell population acts as a trigger point for the cells to differentiate ^5 6^. So scientists and laboratory technicians must harvest the cells before they differentiate and lose their biological potency. Hence, scientists and technicians monitor the growth process by microscopic imaging to ensure quality and to optimize yield.

The microscopic images are often analyzed manually. Hereby, the scientists estimate the fraction of area covered by cells, called confluence. The erroneous process could lead to reduced yield and thus increased cost. We need two things to automatically estimate the confluence: (a) a method to detect and segment cells in images and (b) data from microscopy images annotated with known confluence. Traditional image processing techniques include thresholding methods ^7^ as edge, detection, and region-growing approaches ^8^. However, advances in artificial intelligence (AI) have shown that AI models can outperform traditional methods for cell segmentation ^9^. A typical way to train AI models is to collect data, label it, and train models from scratch i.e. a U-Net model for segmentation ^10^. The use of pre-trained models has recently gained popularity ^11^. Large pre-trained models performing well on various tasks are called foundational models. There are foundational models for computer vision (SAM ^12^, Detectron2^13^) and pre-trained models for cell segmentation (Cellpose ^14^, LiveCell ^15^). These models show promising results in comparison to traditional methods ^9^. While training custom models has also become accessible to end-users ^16^, the advantage of pre-trained models is that they can be used with little to no labeled images. This is especially important since human labeling is costly and time-consuming. Moreover, our specific use case underlines this advantage: MSCs are non-round and irregularly shaped, and ATMP manufacturing processes prohibit staining for higher contrast. Consequently, generating a sufficiently detailed and diverse training dataset for custom model training would require significant effort, further emphasizing the utility of pre-trained models With both (foundation models and untrained models) approaches existing, and labeling being costly, naturally, the question arises of how to estimate confluence with as few labeled data as possible. The amount of data to use ranges from zero-shot learning, where no labeled data is used, to using all available data, with selecting the *n* most informative samples for labeling with active learning (AL) lying in between. AL is a smart way to select the next datapoint/s to label ^17^. There are three main types of active learning: uncertainty-based, diversity-based, and cluster-based ^17^. Using AL to select only a core set of the whole data can produce similar or even better results than labeling all data ^18^, while reducing labeling costs. For our model-driven approach, we focus on uncertainty-based methods.

Since there exists plenty of research for cell segmentation ^9^ and AL ^19 17^, we focus on the intricacies of applying AL to custom small datasets using state-of-the-art models for confluence estimation. In particular, we are interested in four insights: First, we compare the training and fine-tuning performance of four different cell segmentation models on four datasets during the AL dataset curation process. We selected the commonly used models U-Net ^10^, Detectron2^13^, Cellpose ^14^, and Meta’s Segment Anything Model (SAM) ^12^ for our analysis. Specifically, if the dataset curation with AL outperforms a random curation process. Second, we investigate if goal-dependent (confluence estimation) labeling strategies can save dataset curation time/cost while keeping performance. Third, we are interested in how AL selects data points in a microscopy movie context. Fourth, we show if fine-tuning offers a benefit over zero-shot estimation.

## MATERIAL AND METHODS

### Data

Our analysis utilized four distinct datasets, including one external dataset and three from our lab (“internal”). One of these internal datasets contains livecell (lc) imaging data obtained using a CytoSmart Lux microscope (10x magnification; 5-megapixel camera), referred to here as “lc-internal.” We derived a second dataset from the lc-internal dataset by applying different annotation techniques. While, in the original lc-internal dataset each cell was labeled individually, in the second set, cohesive clusters of cells were labeled as single objects. This second dataset is referred to as “lc-internal-lazy”. The images had a dimension of 1280×960 pixels. The goal was to assess whether this simplified annotation was sufficient for confluence prediction. The third dataset includes standard microscopy (“sc-internal”) images taken using a ZEISS Axiovert 40 CFL microscope (10x objective Ph1 (phase contrast); Axiocam ERc 5s, 5-megapixel camera). The images in this dataset were 512×512 pixels in size. We incorporated a larger external dataset from the LIVECell study as a fourth dataset ^15^. To roughly match the cell types of the other datasets, we filtered the external dataset and included only the A172 cell line (which has morphologic similarities to MSC cells despite its origin of glioblastomas). We will refer to this dataset as “lc-external”. Table 1 shows an overview of our dataset, including the number of regions of interest (ROIs), number of images, and image dimensions.

**Table 1:** Dataset characteristics showing the number of regions of interest (ROIs), number of images, and image dimensions for each subset.

| Dataset | #ROIs | #Images | Image Size |
| --- | --- | --- | --- |
| lc-internal-train | 2,243 | 16 | 1280x960 |
| lc-internal-test | 400 | 2 | 1280x960 |
| lc-internal-train-lazy | 897 | 16 | 1280x960 |
| lc-internal-test-lazy | 224 | 4 | 1280x960 |
| lc-external-train | 70,758 | 333 | 702x520 |
| lc-external-test | 30,005 | 138 | 702x520 |
| sc-train | 2,467 | 12 | 512x512 |
| sc-test | 446 | 3 | 512x512 |

We annotated the three internal datasets with the tool ImgLab ^20^ and obtained the annotations in the COCO JSON format ^21^. We obtained annotation instructions from wet lab scientists. We transformed the COCO file to masks with custom scripts (https://git.informatik.uni-leipzig.de/joas/confluence//blob/main/utils/coco_to_mask.py?ref_type=heads) because U-Net and Cellpose require instance masks, instead of COCO JSON files. We split each dataset into a training and test set. Since we did not aim to improve model performances by hyperparameter tuning, we abstained from using a validation split. We used a combination of internal and external datasets, as summarized in Table 1. The lc-external dataset, drawn from the LIVECell paper ^15^, is the largest, with the most ROIs and images in both its training and test splits. The lc-internal dataset includes full and “lazy” subsets for training and testing, with consistent image counts across these variations. Finally, the sc-internal dataset provides additional training and test data with smaller images than the lc datasets. Figure 1 shows sample images and annotations of each dataset. Each ROI corresponds to a labeled cell, and image dimensions vary across datasets, with lc-internal images being the largest and sc-internal images the smallest.

**Figure 1:**
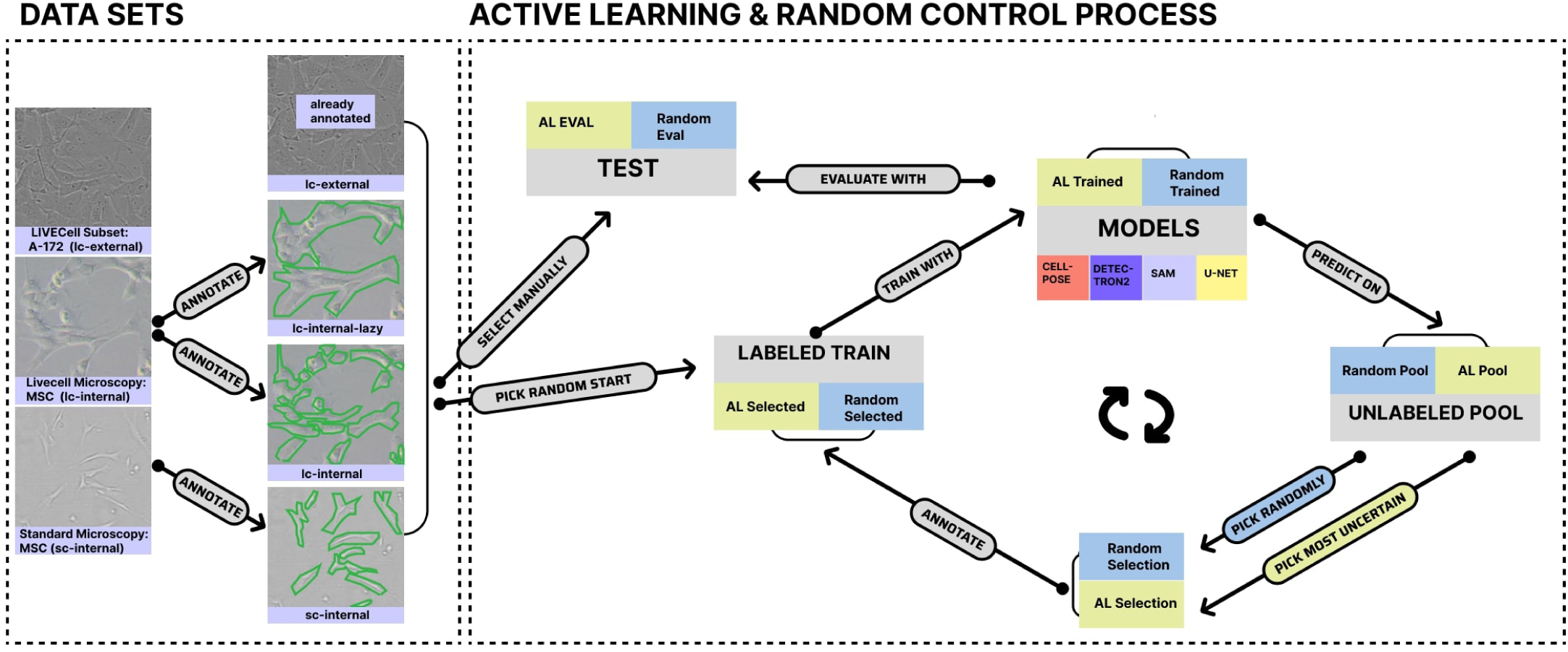
High-level overview of our Active Learning experiment.

### Active Learning for dataset curation

After the dataset curation and processing as described above, we selected the test set manually (for the internal datasets: sc-test, lcinternal-test, and lc-internallazy-test sets), because we wanted to include images with roughly 50% confluence. This confluence range is the most critical in production and thus most important for our models to get right. Next, we defined a pool and a train set. The train set was initially chosen randomly. We selected two to ten images for this initial set, depending on the size of the whole dataset. The selected train images were moved from the pool to the train set (either physically on our machine or by filtering the COCO JSON file). Subsequently, we trained our four models with these initial train sets for 100 epochs. Figure 1 visually represents this process. We trained on NVIDIA A100 GPUs. The exact model configuration and hyperparameters are described in the model section. After training, we evaluated each model on the respective test set. We obtained the predicted instance masks from the model and calculated the image-wise IoU between the ground truth- and predicted masks (intersection between two binary images). We saved the results per image and calculated the mean IoU, standard deviation, and interquartile range across the test set. Additionally, we calculated the absolute differences in the confluence between ground truth and predictions.

In addition to evaluating on the test set, we performed inference on the pool set, generating probability maps for each image (except for the Detectron2 model, as detailed below). We then computed the Shannon entropy of these probability maps and ranked all pool set images based on their entropy values. We move the image with the highest entropy from the pool to the train set (for the lc-external dataset, we move ten images from pool to train). Additionally, we add annotation information by updating the COCO JSON file or generating the instance masks for this image. This approach, known as entropy-based sampling ^17 22^, is a widely used technique in uncertainty-based active learning ^23^. We iteratively retrain the model as the training set expands, repeating the process until all images are transferred from the pool set to the training set. After each training iteration, we evaluate the model on the test set. As a control, we conducted an additional experiment where images were randomly selected from the pool set to be added to the training set. This results in ten iteration steps for the standard microscopy dataset, 14 steps for the lc-internal datasets, and 34 steps for the lc-external dataset.

To account for randomness in the selection of the initial train set and the random control experiment, we repeated the above-described process ten times in total. We aggregated the above-described evaluation metrics at each iteration step over all ten experiments with the mean. Additionally, we calculated the interquartile range of the metrics across the random experiments. To analyze if AL significantly improves training, we compared the random picks with the AL picks and applied the Mann-Whitney U-Test ^24^ at each step for the means of the evaluation metrics across all ten experiments. We ran this experiment with all four models on all four datasets.

### Goal dependent labeling

Most segmentation tasks care about the actual shape of the object. In adherent cell cultures cells often grow together to a blob/cluster. Labeling a blob of cells takes much less effort than each cell separately. When caring only about the area covered by cells for confluence estimation, we hypothesized that is enough to label blobs of cells. Therefore we compared the model performance between the lcinternal and lc-internal-lazy datasets. We performed the same AL process as described in the above section, but instead of comparing AL against random image selection, we compared the evaluation metrics of the two different labeling strategies per step. We used the Mann-Whitney U Test ^24^ significance.

### Active Learning in a Microscopy movie context

The images of the “sc-internal” dataset originated from a microscopy movie. This means the later the images were taken in the movie, the more the cells have grown (higher confluence). Hence, we expect the movie positions to influence the AL selection process. We took the results from the AL process described in the first section and calculated how many positions in the movie the selected image differs from the image selected in the previous step. We again aggregated this by the mean and calculated the interquartile range across all ten random runs. As a control, we compare the image selected by AL with the randomly selected image. We tested with the Mann-Whitney U Test ^24^ for significance.

### Fine-tuning compared to zero-shot learning

In our AL experiment, we evaluated the necessity of fine-tuning by first using all models without additional training and directly feeding the test set into them for inference. At the opposite end of the spectrum, we fine-tuned the models using all available labeled data, training them for 500 epochs (without early stopping). We used the models with the pre-trained weights for the zero-shot experiment but did not fine-tune on our four datasets. We performed the inference on the test set. We used IoU and the absolute delta in confluence as evaluation metrics. To set the performance of the models in a better context, we included a baseline confluence detector that does not incorporate modern deep learning. This baseline algorithm processes grayscale images and detects edges with the Canny edge detector ^25^. It then fills gaps in the detected edges using binary hole-filling. Next, it removes small objects and detects contours in the processed image with the “marching squares” algorithm, finding iso-valued contours at a specific level. After detecting the contours, any open contours are closed and simplified with polygonal approximation. Our baseline algorithm then draws these contours onto a blank mask and interpolates between them to create a filled mask.

## Models

### Cellpose

We used the custom fork of the Cellpose model from Singer et al. ^14^ based on version 3.0.0. We made no modifications to the main model. Instead, we implemented functionality to allow custom names for standard Cellpose log files. As of this writing, this feature has been incorporated into Cellpose based on our pull request. We used the train function from the Cellpose model with the model type “cyto” for fine-tuning. Cellpose takes the diameter mean of the cells as input. We calculated the cell diameter mean with a custom script (https://git.informatik.uni-leipzig.de/joas/confluence/-/blob/main/cellpose_main.py?ref_type=heads) for our data. For inference, Cellpose requires two key thresholds: the cell probability threshold and the flow threshold. The cell probability threshold determines the minimum probability for pixels to be classified as part of a cell, while the flow threshold controls the tolerance for errors in detecting cells ^14^. We observed that the model’s performance is sensitive to these thresholds, so we implemented an automatic tuning process to optimize them for our train set. We designed a method that systematically explores different threshold combinations. The function evaluates the performance of the model on the training set, using ground truth masks as a reference. It iteratively tests a range of flow thresholds (from 0 to 3) and cell probability thresholds (from −6 to 6) in 0.5 steps to identify the optimal combination.

The model generates predicted masks for each threshold pair, which are preprocessed before being compared to the ground truth masks. The number of detected cells is calculated for both, the predicted and the ground truth masks. We optimize the thresholds for IoU score. Additionally, a penalty is applied if no cells are detected in the predicted masks, ensuring that the model does not optimize for precision and only outputs background and no masks. The function selects the combination of thresholds that maximizes the score, which is determined by the IoU with the applied penalty. The thresholds corresponding to the highest score in the train are then chosen as the optimal values for inference. We did not change any other hyperparameters from the default values of the Cellpose model class and the Cellpose train method.

For the AL part, we obtained the cell probability directly from the Cellpose model’s eval method and calculated each pool image’s Shannon entropy. Furthermore, we trained all models in the AL experiments for 100 epochs.

### Detectron2

For our analysis, we employed the Detectron2 framework to perform instance segmentation. The model was set to detect a single class, corresponding to the cells in our images. The exact model configuration can be found in our code repository (https://git.informatik.uni-leipzig.de/joas/confluence/-/blob/main/utils/utils.py) To determine which image to label next in the AL process, we cannot obtain the probability masks directly from Detectron2. Instead, the model provides already calculated confidence scores per mask. We use the average of these scores as a metric of uncertainty. In the inference step, we select the image(s) with the lowest score to label next.

### Segment Anything Model

We extended SAM, which does not natively support fine-tuning, by constructing a custom module to enable this functionality. We published a blogpost (https://maxjoas.medium.com/finetune-segment-anything-sam-for-images-with-multiple-masks-34514ee811bb) that makes fine-tuning SAM publicly accessible. Our implementation is based on SAM version 1.0.0^12^. We fine-tuned the model by creating a custom wrapper around its architecture, allowing training on a new dataset. Specifically, we used the PyTorch *nn.module* class, where we selectively froze the image encoder and prompt encoder for computational efficiency, while leaving the mask decoder trainable. Since we do not know the spatial location of the cells beforehand, we do not provide bounding boxes as prompts. We resized the input images to 1024×1024, the same dimensions SAM used during its original training. We loaded the model using the *vit_h_* variant from SAM’s model registry and initialized it with this pre-trained checkpoint. During the forward pass, the model generated low-resolution probability mask maps, which we upsampled to match the original image size. We used a combination of Focal and Dice loss with a weighting of 20:1 as in the SAM paper ^12^ For the AL process, we calculated the Shannon Entropy of the predicted probability maps and selected the image(s) with the highest entropy to move from the pool to the train set.

### U-Net

We used U-Net based on the original architecture proposed by Ronneberg et al. ^10^. We describe the training process and hyperparameters in our GitLab repository (https://git.informatik.uni-leipzig.de/joas/confluence-unet)

We obtained the mask probability map directly from the U-Net model for the AL process. We calculated the Shannon Entropy of the predicted probability maps and selected the image(s) with the highest entropy to move from the pool to the train set.

## RESULTS

### Active Learning Performance

We analyzed if AL is an approach to improve cell segmentation and to reduce labeling effort when using four commonly used models for segmentation. We show that uncertainty-based AL offers no improvement in dataset curation for confluence prediction in our use case. The experiments with Cellpose and SAM showed the least differences between random and AL image selection with only four (Cellpose) and three (SAM) significantly different steps out of 72 total steps (cf. Figure 2 and Supplement Figure S1 for IoU). We observe a significantly better performance for the U-Net model with random dataset curation for the external dataset. Detectron2 is the only model where AL improved confluence prediction in the standard microscopy dataset and to some extent in the external dataset.

**Figure 2:**
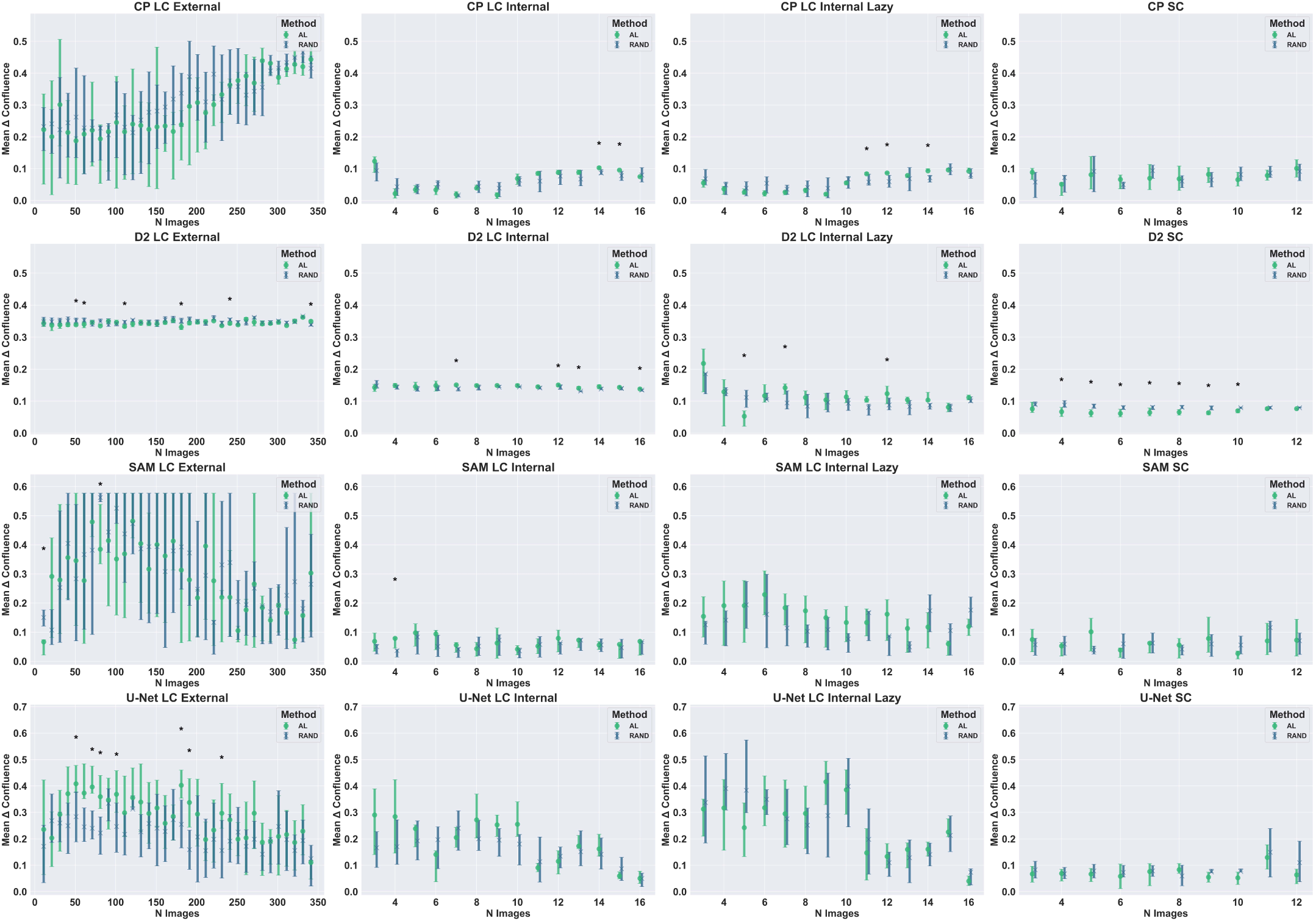
Impact of active learning on dataset curation. Each plot represents one model-dataset combination. It shows the mean difference between the true and predicted confluence across the ten experiments at each step. One step represents the addition of newly labeled images selected randomly (blue) or by AL (green) The error bars show the interquartile range. Significant differences (Mann-Whitney U-Test) are marked with an asterisk (p-value *<* 0.05). Abbreviations: CP: Cellpose, D2: Detectron2

We hypothesized that specialized and non-specialized pre-trained segmentation models benefit from fine-tuning since the shape and image characteristics of MSC cells are unique and complex. However, the effect of fine-tuning is limited and largely dependent on the dataset. In seven experiments, we observed the best scores during the early stages of fine-tuning in AL, showing that more data does not always improve results. Our control experiments with random picks for the next image to label showed similar results with six of the best performances coming from the first half of the fine-tuning. From a model perspective, we observe that Detectron2 does not benefit from fine-tuning, Cellpose gets even worse, and for SAM and U-Net, we do not see a clear trend.

In particular, nine out of 16 experiments (model and dataset combinations) achieved a minimal delta in the confluence of not more than 0.05, and three experiments achieved a minimal delta in the confluence of no more than 0.10. Comparing the mean of performances across all datasets, SAM predicts the confluence the most exactly with a value of 0.05 *±* 0.02 and Detectron2 0.15*±* 0.13. Performance analysis across datasets reveals that the lazily labeled dataset achieved the best results (Mean 0.04 *±* 0.02), whereas the external dataset had the worst performance (Mean 0.18 *±* 0.11). Table 2 shows the aggregation of the best results across models and datasets in detail. Non-active learning (randomized) shows similar trends (see Table S1). Table S2 shows the delta confluence metrics for all experiments. We observe similar trends when using IoU as performance metrics as shown in Table S4 and Table S5.

**Table 2:** Mean and standard deviation of minimum delta confluence values across models and datasets. The column *Convergence Point* represents the fraction of Active Learning iterations completed, where 0 indicates no additional data added, and 1 signifies that all available data has been included.

| Model | Min | Convergence Point |
| --- | --- | --- |
| Cellpose | $0.07 \pm 0.08$ | $0.29 \pm 0.20$ |
| Detectron2 | $0.15 \pm 0.13$ | $0.50 \pm 0.37$ |
| SAM | $0.05 \pm 0.02$ | $0.56 \pm 0.41$ |
| U-Net | $0.06 \pm 0.03$ | $0.95 \pm 0.11$ |

(b) Dataset Aggregation (See Data section for more details)
| Dataset | Min | Convergence Point |
| --- | --- | --- |
| lc-external | $0.18 \pm 0.11$ | $0.41 \pm 0.45$ |
| lc-internal | $0.06 \pm 0.05$ | $0.75 \pm 0.29$ |
| lc-internallazy | $0.04 \pm 0.02$ | $0.63 \pm 0.40$ |
| sc-internal | $0.05 \pm 0.01$ | $0.50 \pm 0.34$ |

### Goal-dependent labeling

For confluence prediction, precise segmentation of individual cells is unnecessary. We investigated whether faster labeling of cell clusters as single units (“lazy labeling”) impacts model performance. The results achieved through goal-specific labeling do not universally enhance every model’s performance. We see a clear trend in the Detectron2 model with ten out of 14 steps being significantly better for the lazy labeling method. Whereas for U-Net and Cellpose we can see no clear difference. The SAM model even shows better results for the exact labeled images concerning confluence estimation. However, the IoU is significantly higher by a large margin when labeling the data lazy for all models. Figure 3 shows the comparison of the two labeling methods with IoU and delta confluence for all four models.

**Figure 3:**
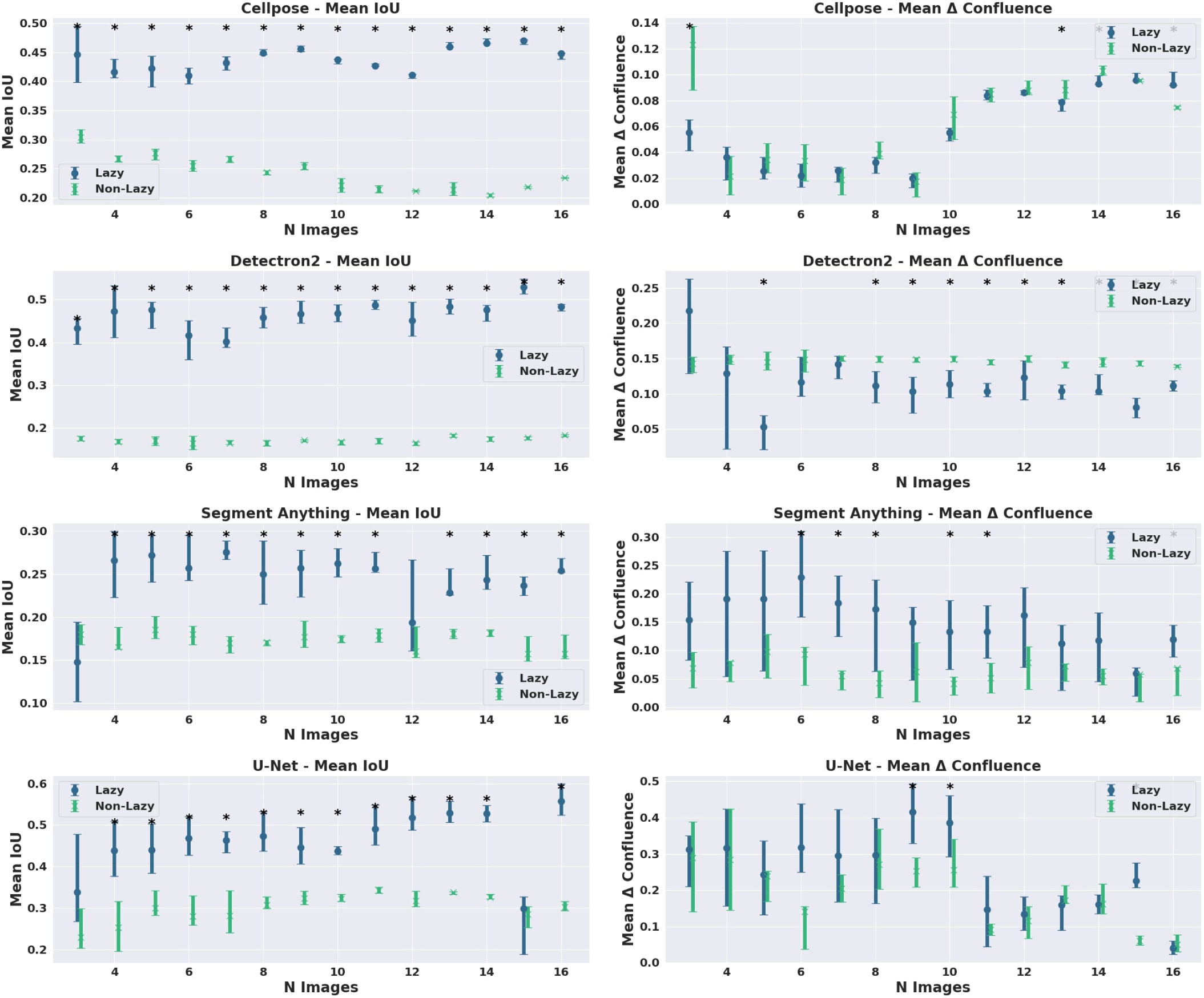
Impact of lazy labeling. The plot shows the performance comparison of lazy and exact labeling methods during the dataset curation process for all models. The first column shows the IoU metric and the second column the differences in confluence.

### Active Learning in a Microscopy movie context

We expected that uncertainty-based AL preferably selects images with varying confluence which sequentially increases in our standard microscopy dataset recorded as a movie of a cell culture. Overall, we observe no consistent difference or tendency in the movie position of the selected images between the AL and random selection process across our four segmentation models (see Figure 4). This indicates that the cell density is no factor in the model’s uncertainty, likely, because cell shape is not heavily altered over time. Further, results are in line with the observation that AL does not significantly improve model performance for cell segmentation and confluence prediction as shown in Figure 2.

**Figure 4:**
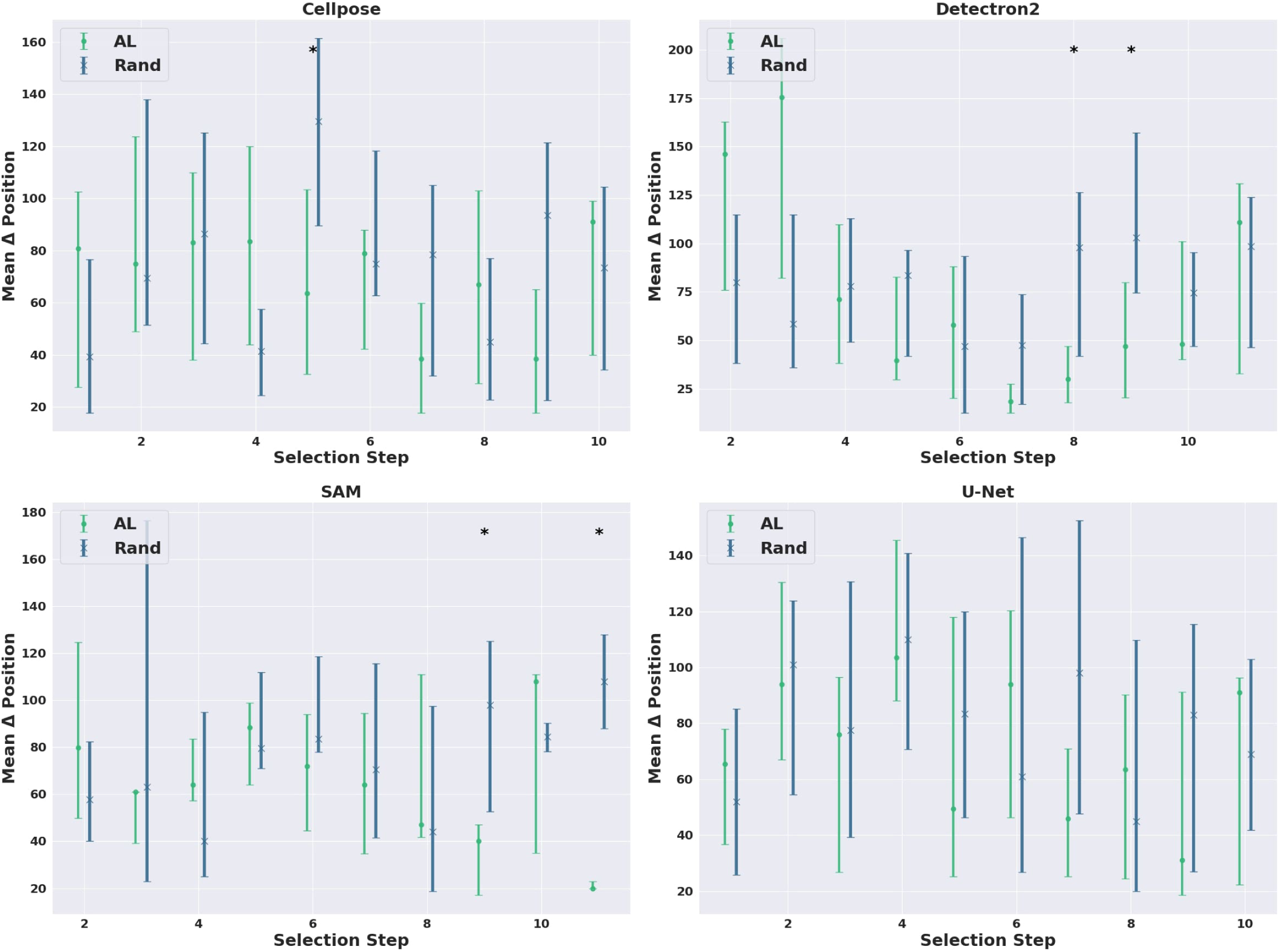
Selection pattern of active learning. Each subplot shows the mean difference in movie position between the selected and the previously selected image at each step for the AL and random selection for each model.

### Zero-shot inference or full fine-tuning

Since we observed that models showed mixed behavior during fine-tuning e.g. Cellpose showing a decrease in performance, we analyzed more deeply the zero-shot capability of cell segmentation models. The performance of SAM is with a delta of 0.05 *±*0.036 in confluence estimation near perfect across all datasets even without fine-tuning. Deep learning-based approaches outperform traditional image analysis techniques by a large margin. We see a substantial performance increase for Detectron2 when applying fine-tuning. However, this is dependent on the dataset, since the performance on the external data decreased. We see small dataset-dependent increases for the Cellpose model. Figure 5 shows the difference in true and predicted confluences for all model-dataset combinations. In summary, our results show a strong indication that for confluence prediction (for MSC-like cells) generalist foundation models like SAM outperform specialized (foundation) models (Cellpose), and fine-tuning is not necessary. Further, in the case of Cellpose results indicate that fine-tuning with irregular cell shapes (MSC) can lead to decreasing performance rather than expected improvements.

**Figure 5:**
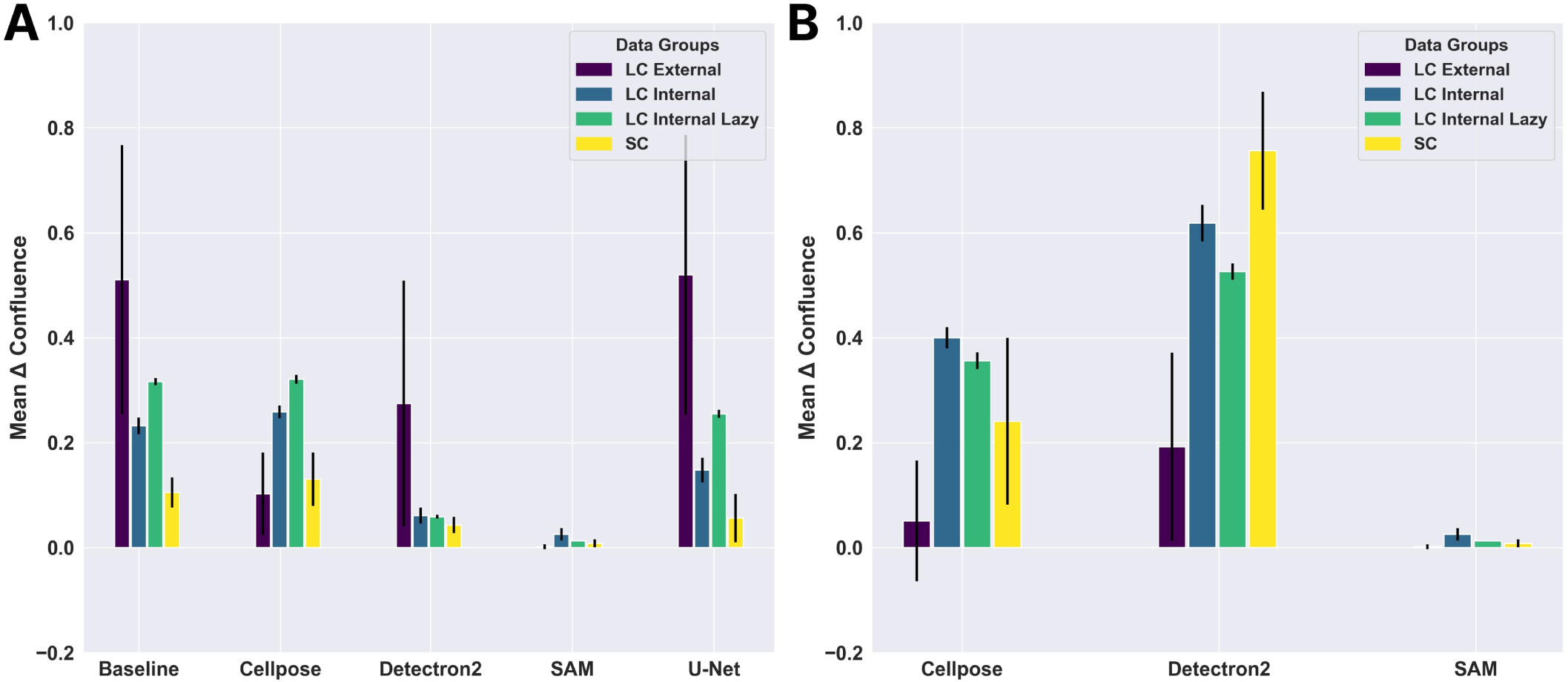
Comparison of fine-tuning and zero-shot results per model and dataset. Figure A shows the differences in true and predicted confluence for four models and one baseline across all datasets for fine-tuning. Figure B shows the performance of zero-shot learning for models where zero-shot is applicable.

### Qualitative Analysis / Usability

When deciding how to curate a dataset for confluence estimation and which model to choose, there are many practical considerations besides pure performance. We will give insights on (a) the difficulty of fine-tuning the models (b) implementing an uncertainty-based AL approach and (c) on overall computational considerations.

Detectron2^13^ is the easiest model to fine-tune since fine-tuning is a built-in capability. The documentation for Detectron2 is clear and easy to follow. Detectron2 requires the input data as COCO annotations, which is a widely used format. The ease of use comes with less customizability and control. Detectron2 is not the most cutting-edge model and does not specialize in cell segmentation. Cellpose ^14^ specializes in cell segmentation and offers fine-tuning options. However, Cellpose predictions are highly influenced by cell probability and flow thresholds. These thresholds can be adjusted when knowing the ground truth, but for automatic segmentation on unknown data, this is not possible and would require manual intervention to find the best thresholds. Cellpose requires ground truth masks as input for fine-tuning, which is also customary. With releases in February 2024, Cellpose offers up-to-date models and some customizability regarding the cell type.

SAM ^12^ was the hardest model to fine-tune because this option is unsupported. We needed to write custom wrapper classes to enable fine-tuning. This is not possible without major technical expertise in deep learning. On the other hand, SAM is easy to use for zero-shot learning and is the most potent of all used models. SAM requires the annotations in the common COCO JSON format.

We did train U-Net from scratch and used no fine-tuning. U-Net requires implementation knowledge, e.g. Pytorch or Keras, and training from scratch. Due to its limited performance, it does not provide a good trade-off between efficient confluence prediction.

To curate a dataset with AL we need to obtain uncertainty measures from the models. Detectron2 was the only model that returned confidence scores for the mask predictions. However, there was no built-in functionality to obtain probability masks. In our custom implementations of U-Net and SAM, we made it easy to obtain the probability maps directly. Nevertheless, this precedes custom implementation which is hard to do in some cases. Cellpose returns the cell probabilities directly and thus an uncertainty-based AL approach is straightforward to implement.

We recommend a GPU for model training or fine-tuning for all four models. For inference, a CPU suffices for U-Net, Detectron2, and Cellpose. This is particularly advantageous for integrating confluence prediction into real-world automation systems for cell production, where high-performance GPUs may not be readily available. However, inference with SAM on a CPU is not practical due to the model’s size and slow performance.

The fine-tuning process requires a GPU for all models to ensure completion within a reasonable time. Fine-tuning on larger datasets using a CPU is impractical, as it could take weeks and offers minimal benefit compared to zero-shot training. Conversely, fine-tuning with only a few images can be completed in under a week (and even within hours for Detectron2 and Cellpose) and provides notable performance improvements for these models. Due to the high computational cost and lack of significant performance gains, we do not recommend fine-tuning SAM.

## DISCUSSION

We could not observe the expected advantages of AL in dataset curation. We attribute this to (a) limited diversity within the dataset, (b) mainly use of large pre-trained models, (c) a primitive AL approach, and (d) the simple binary classification (fore/background) task. Firstly, since our datasets only contain one cell type, the primary variation lies in the growth state or cell density. This does not fundamentally alter the characteristics of the objects to be segmented (i.e., the cells). Consequently, the timing of when a given example is presented during training has a limited impact on model performance. Our analysis revealed that AL did not preferentially select images based on their temporal position in the growth sequence, showing no significant difference from random selection. This finding supports our explanation that cell density alone does not create the kind of meaningful variation that AL strategies typically exploit ^17^. The model’s uncertainty, which drives the AL selection process, appears to be independent of the growth state, suggesting that once the model learns to segment cells at one density, it can readily generalize to other densities.

Secondly, we hypothesize that the benefit of AL in pre-trained models is minimal due to the extensive data exposure these models have already undergone, which reduces the impact of new data points. Additionally, our observations suggest that U-Nets struggle when presented with a small number of highly diverse instances through Active Learning, whereas random selection maintains a distribution more representative of the complete dataset, allowing the U-Net to properly specialize. Thirdly, for usability reasons, we employ a straightforward maximum entropy approach for uncertainty-based AL. However, capturing the complexities of this data and model may require more sophisticated AL strategies, often involving combinations of techniques designed to handle nuanced variations more effectively. While prior research on biomedical images shows that various AL techniques (including uncertainty-based ones) can achieve comparable performance with a reduced number of data samples ^26 27 28 29^, these did not include pre-trained models, used more diverse data or more complex AL algorithms on top of an uncertainty-based approach.

We wanted to investigate whether a simple AL approach could help with dataset curation and our finding emphasizes the need for careful assessment of AL’s utility in each unique scenario. AL should be best used when pre-trained models are not suitable for a use case and sophisticated AL algorithms are accessible for one’s specific problem.

Beyond active learning, another effective approach to dataset curation is tailoring the labeling strategy directly to the desired outcome i.e. confluence estimation. For the IoU metric, lazily labeled data consistently outperformed traditional labeling approaches. We attribute this to simpler, shapes that are easier for models to learn. Even when considering the confluence task directly, instead of IoU, we observe no drop in performance when labeling lazily. This is likely because precise annotation does not negatively impact confluence estimation, yet achieving accurate confluence estimation does not require highly precise labeling. Both approaches deliver satisfactory confluence estimation, making the differences between them less significant for this metric. Interestingly, when using the SAM model, traditionally labeled data performed slightly better, which we attribute to SAM’s extensive pre-training on precisely annotated datasets, making it less effective at leveraging lazily labeled examples. Alternatively, lazy labeling could introduce irreducible error and act as a source of noise. Despite the less clear advantages in terms of delta confluence accuracy, the substantial reduction in labeling effort makes the lazy labeling approach attractive for confluence estimation. In particular, since detailed cell segmentation is not required (e.g. to derive individual cell characteristics), a simple fore-background classification is in this use-case. While this specific labeling strategy may not generalize to all problems, it demonstrates the value of developing task-specific labeling approaches that balance annotation effort with model performance.

To our knowledge, no existing research directly addresses confluence estimation-specific annotation. However, recent studies in biomedical imaging have highlighted the use of time-efficient annotation techniques combined with self- and semi-supervised learning to reduce labeling demands while maintaining high performance. For example, in nuclei segmentation, some approaches focus on selectively annotating only a small subset of critical image patches, using semi-supervised methods and data augmentation to match fully supervised model performance while minimizing labeled data requirements ^30^. Similarly, in cell segmentation, weakly supervised methods use single-point annotations per cell, combined with self- and co-training strategies, to achieve segmentation accuracy close to that of fully supervised methods with substantially less annotation effort ^31^. Krishnan et al.’s review further highlights how efficient annotation methods, when used with self-supervised learning, allow models to capitalize on large volumes of unannotated data, enhancing model development while reducing expert annotation time ^32^. Furthermore, tools like LABKIT ^33^ provide interfaces for efficient annotation, combined with supervised deep learning. Although we implemented only a time-efficient annotation strategy without any form of weak supervision, given the relative simplicity of our task, these findings still support our results. They suggest that efficient annotation methods, especially when combined with self- and semi-supervised learning, can significantly reduce labeling effort in active learning while maintaining model performance.

While efficient labeling strategies can reduce annotation effort, eliminating the need for labeling would even more practical. Our experiments with zero-shot inference demonstrated that for simple tasks like confluence estimation, the largest foundation model available, SAM, achieved nearly perfect confluence estimation without any task-specific training. This indicates that the cost-benefit ratio of fine-tuning declines the more advanced the models get. While other models showed marginal improvements with fine-tuning, and even SAM demonstrated slight gains, these benefits were minimal compared to SAM’s excellent zero-shot performance. Interestingly, across all models, we found that the best performance was achieved using only a subset of the available training data, suggesting that more data does not necessarily lead to better results. The costs associated with fine-tuning—including substantial computational resources, technical expertise for model adaptation, and time investment in data collection and annotation—far outweigh these modest performance gains. This cost-benefit analysis strongly favors using SAM in its zero-shot configuration for basic confluence estimation tasks, eliminating the need for resource-intensive fine-tuning. Further, the recently released SAM 2^34^ may show even higher accuracy for such tasks and has a focus on object tracking in video contexts which is of high interest in time-series data from cellular production systems. When comparing our results with existing research, we find that zero-shot inference with SAM is powerful even for more complex tasks. However, the optimal approach appears task-dependent and some amount of fine-tuning or combining it with other models still seems to have a benefit: Baral and Paing ^35^ used a fine-tuned object detection model (YOLOv9-E) to generate prompts for zero-shot SAM inference, followed by traditional image processing refinements. This hybrid approach achieved highly accurate performance (>94% mAP50) for cell segmentation across varying difficulty levels, without fine-tuning SAM directly. In contrast, more specialized applications may warrant fuller investment in model adaptation. CryoSeg-Net ^36^ demonstrated that combining SAM with a task-specific U-Net significantly improved protein particle detection in cryo-EM images compared to using SAM alone, but this required extensive dataset curation and model engineering. Similarly, the Segment Anything for Microscopy project ^37^ showed that specialized training for multi-dimensional microscopy data substantially enhanced segmentation quality across diverse imaging conditions, justifying the additional investment due to the complexity of volumetric segmentation and tracking tasks. These varying approaches highlight the importance of carefully considering task complexity and resource constraints when choosing between zero-shot applications, hybrid solutions, and full fine-tuning of foundation models.

While our findings provide insights into confluence estimation for MSC production standardization, we acknowledge several limitations of our study. First, our results are specifically focused on confluence estimation, reflecting the practical needs of our project-based research. However, the principles we uncovered regarding model selection and data curation strategies may inform similar biomedical image analysis tasks. Our model selections, while not exhaustive, strategically covered a representative spectrum of approaches: from models trained from scratch to cell-specific models and powerful general-purpose foundation models. This selection allowed us to compare different paradigms in model development while maintaining practical feasibility. Similarly, while our datasets were limited to one cell type, they enabled us to draw important conclusions about the influence of data diversity on active learning effectiveness. Despite these limitations, our study makes several valuable contributions: We demonstrated that for homogeneous cell cultures, SAM delivers excellent results without the need for resource-intensive approaches such as active learning or fine-tuning. We provide technical guidelines for implementing active learning in cell imaging applications, demonstrate the importance of goal-specific labeling strategies, and highlight how data homogeneity affects active learning performance. These insights can guide future research in automated cell culture monitoring and quality control. These practical insights significantly lower the barrier to implementing automated quality control in cell manufacturing. A prototype of our confluence detection software is publicly (https://livinglab.scadsai.uni-leipzig.de/cell-confluence/), enabling immediate community adoption.

## AUTHOR CONTRIBUTIONS STATEMENT

## ACKNOWLEDGMENTS

MJ and DF acknowledge support by the German Federal Ministry of Education and Research under the funding code 03ZU1111NC and 03ZU1111NB (SaxoCellSystems) as part of the Clusters4Future cluster SaxoCell. RH and JE acknowledge the financial support by the Federal Ministry of Education and Research of Germany and by Sächsische Staatsministerium für Wissenschaft, Kultur und Tourismus in the programme Center of Excellence for AI-research “Center for Scalable Data Analytics and Artificial Intelligence Dresden/Leipzig”, project identification number: ScaDS.AI The authors gratefully acknowledge the computing time made available to them on the high-performance computer at the NHR Center of TU Dresden. This center is jointly supported by the Federal Ministry of Education and Research and the state governments participating in the NHR (www.nhr-verein.de/unsere-partner).

## SUPPLEMENTARY INFORMATION

### Active learning vs random selection with IoU metric

**Figure S1:**
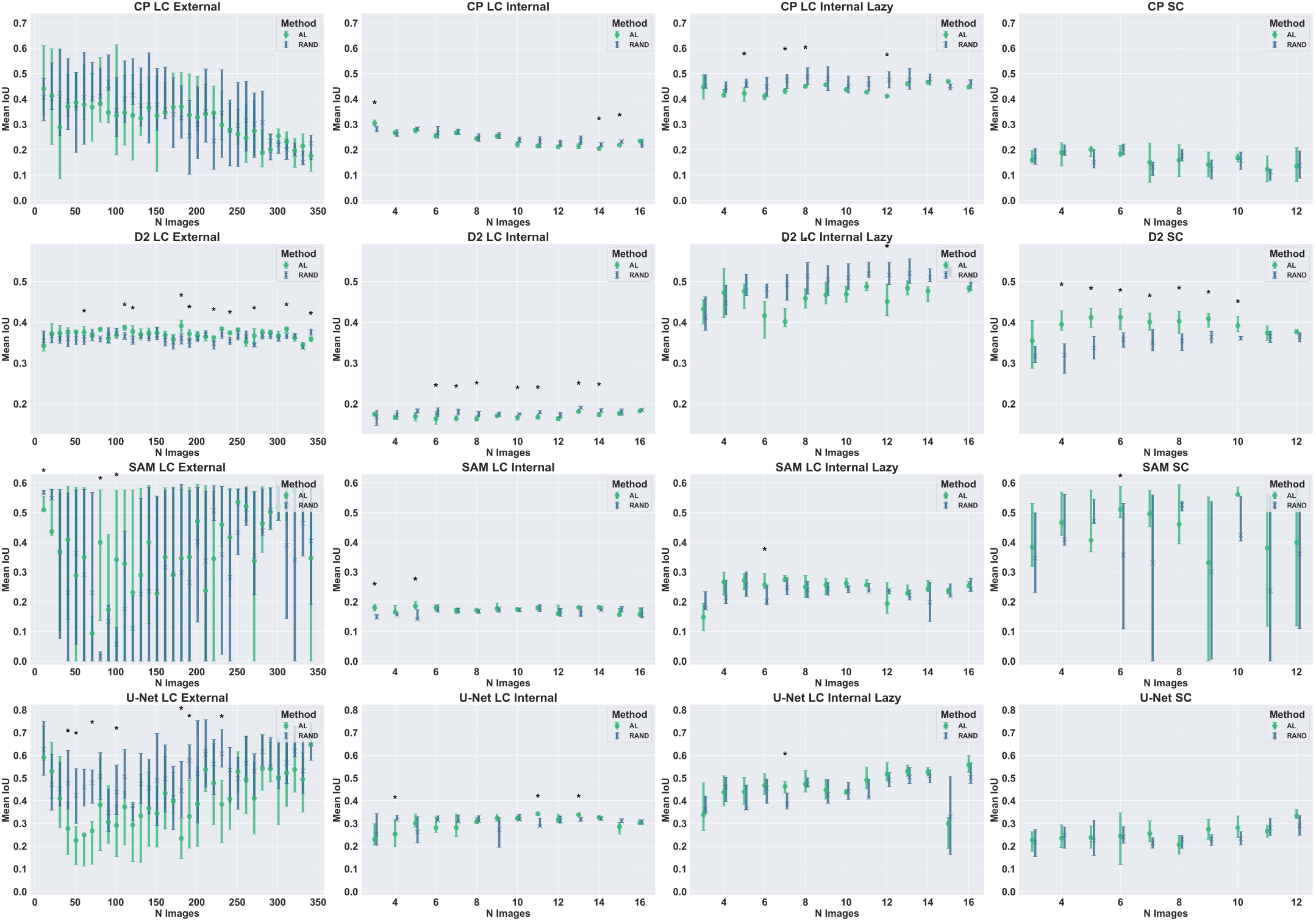
Each plot represents one model dataset combination. It shows the mean IoU for the ten experiments at each step. One step represents the addition of newly labeled images selected randomly (blue) or by AL (green) The errorbars show the interquartile range. Significant differences are marked with an asterisk.

### Overview of best performances for the random dataset curation experiment

**Table S1:**
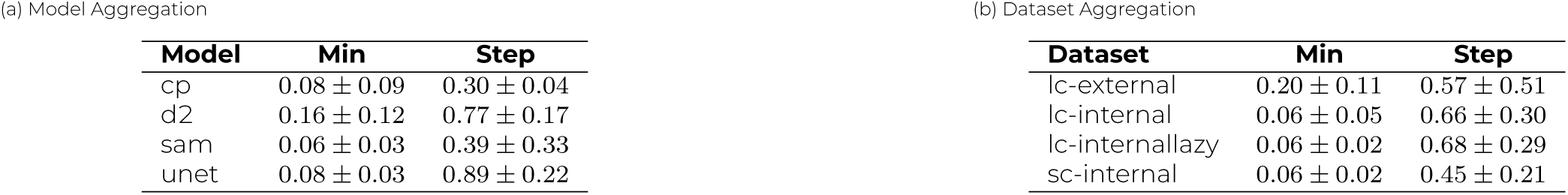
Mean and standard deviation of minimum values and best relative step across models and datasets.

(a) Model Aggregation
| Model | Min | Step |
| --- | --- | --- |
| cp | $0.08 \pm 0.09$ | $0.30 \pm 0.04$ |
| d2 | $0.16 \pm 0.12$ | $0.77 \pm 0.17$ |
| sam | $0.06 \pm 0.03$ | $0.39 \pm 0.33$ |
| unet | $0.08 \pm 0.03$ | $0.89 \pm 0.22$ |

(b) Dataset Aggregation
| Dataset | Min | Step |
| --- | --- | --- |
| lc-external | $0.20 \pm 0.11$ | $0.57 \pm 0.51$ |
| lc-internal | $0.06 \pm 0.05$ | $0.66 \pm 0.30$ |
| lc-internallazy | $0.06 \pm 0.02$ | $0.68 \pm 0.29$ |
| sc-internal | $0.06 \pm 0.02$ | $0.45 \pm 0.21$ |

### Oveview of confluence prediction performances of all experiments in the active learning dataset curation process

**Table S2:** Best Delta Confluence Performance Metrics for Active Learning Experiments.

| Model-Dataset | Min | Max | Best Step | Convergence Point |
| --- | --- | --- | --- | --- |
| unet-lc-internal | 0.05 | 0.29 | 13.0 | 1.0 |
| d2-lc-internal | 0.14 | 0.15 | 13.0 | 1.0 |
| sam-lc-internal | 0.04 | 0.1 | 7.0 | 0.54 |
| cp-lc-internal | 0.02 | 0.12 | 6.0 | 0.46 |
| unet-lc-external | 0.11 | 0.41 | 330.0 | 1.0 |
| d2-lc-external | 0.33 | 0.36 | 170.0 | 0.52 |
| sam-lc-external | 0.07 | 0.48 | 0.0 | 0.0 |
| cp-lc-external | 0.19 | 0.44 | 40.0 | 0.12 |
| unet-sc-internal | 0.05 | 0.13 | 7.0 | 0.78 |
| d2-scinternal | 0.06 | 0.08 | 3.0 | 0.33 |
| sam-scinternal | 0.03 | 0.1 | 7.0 | 0.79 |
| cp-scinternal | 0.05 | 0.1 | 1.0 | 0.11 |
| unet-lc-internallazy | 0.04 | 0.42 | 13.0 | 1.0 |
| d2-lc-internallazy | 0.05 | 0.22 | 2.0 | 0.15 |
| sam-lc-internallazy | 0.06 | 0.23 | 12.0 | 0.92 |
| cp-lc-internallazy | 0.02 | 0.1 | 6.0 | 0.46 |

### Oveview of performances of all experiments in the random dataset curation process

**Table S3:**
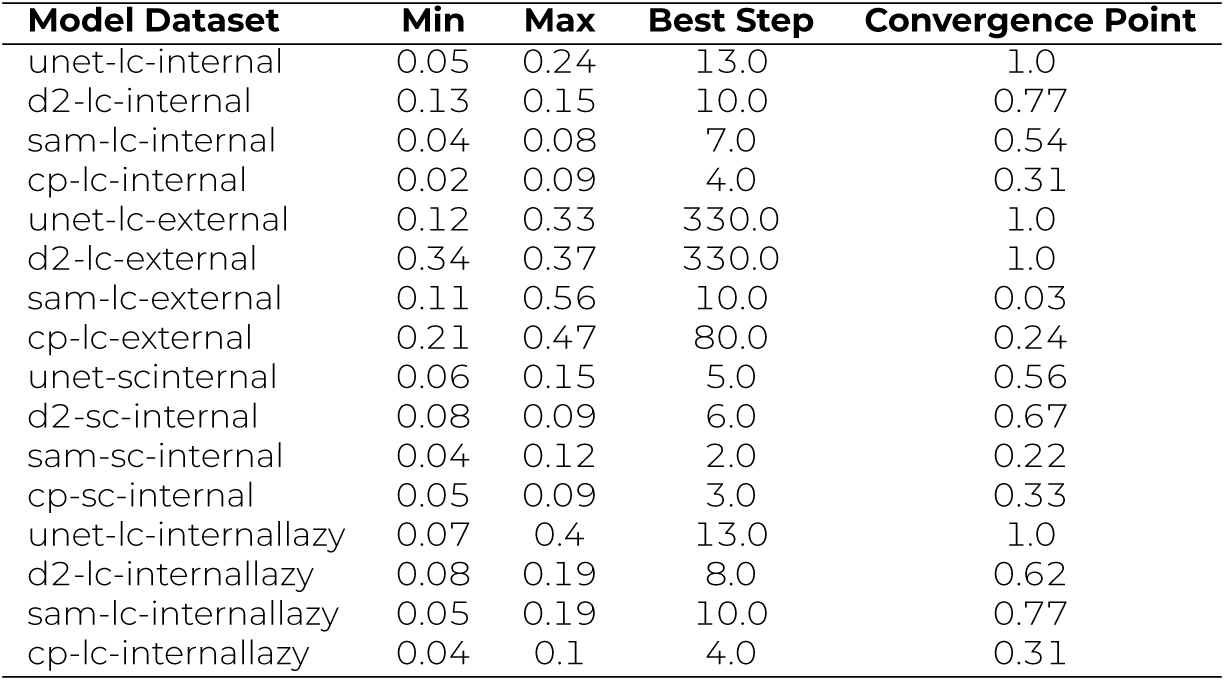
Best Confluence Performance Metrics for Random Experiments.

### Oveview of IoU prediction performances of all experiments in the active learning dataset curation process

**Table S4:** Best Intersection over Union Performance Metrics for Active Learning Experiments.

| **Model Dataset** | **Min** | **Max** | **Best Step** | **Convergence Point** |
| --- | --- | --- | --- | --- |
| unet-lc-internal | 0.23 | 0.34 | 8.0 | 0.62 |
| d2-lc-internal | 0.16 | 0.18 | 13.0 | 1.0 |
| sam-lc-internal | 0.16 | 0.19 | 2.0 | 0.15 |
| cp-lc-internal | 0.2 | 0.3 | 0.0 | 0.0 |
| unet-lc-external | 0.23 | 0.65 | 330.0 | 1.0 |
| d2-lc-external | 0.34 | 0.39 | 170.0 | 0.52 |
| sam-lc-external | 0.09 | 0.58 | 310.0 | 0.94 |
| cp-lc-external | 0.18 | 0.44 | 0.0 | 0.0 |
| unet-sc-internal | 0.21 | 0.33 | 9.0 | 1.0 |
| d2-scinternal | 0.35 | 0.41 | 3.0 | 0.33 |
| sam-sc-internal | 0.33 | 0.56 | 7.0 | 0.78 |
| cp-sc-internal | 0.12 | 0.2 | 2.0 | 0.22 |
| unet-lc-internallazy | 0.3 | 0.56 | 13.0 | 1.0 |
| d2-lc-internallazy | 0.4 | 0.53 | 12.0 | 0.92 |
| sam-lc-internallazy | 0.15 | 0.28 | 4.0 | 0.31 |
| cp-lc-internallazy | 0.41 | 0.47 | 12.0 | 0.92 |

### Oveview of IoU prediction performances of all experiments in the random dataset curation process

**Table S5:**
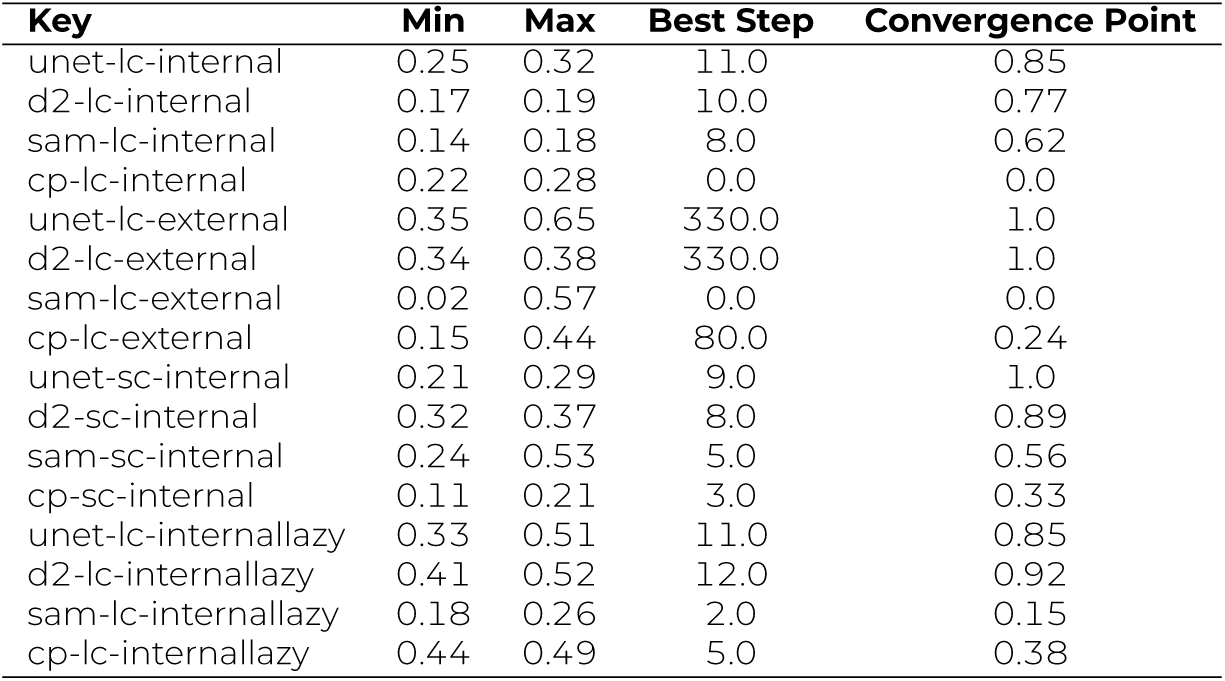
Results across model-dataset combinations for the random experiment with minimum and maximum values, best step, and best relative step.

| Key | Min | Max | Best Step | Convergence Point |
| --- | --- | --- | --- | --- |
| unet-lc-internal | 0.25 | 0.32 | 11.0 | 0.85 |
| d2-lc-internal | 0.17 | 0.19 | 10.0 | 0.77 |
| sam-lc-internal | 0.14 | 0.18 | 8.0 | 0.62 |
| cp-lc-internal | 0.22 | 0.28 | 0.0 | 0.0 |
| unet-lc-external | 0.35 | 0.65 | 330.0 | 1.0 |
| d2-lc-external | 0.34 | 0.38 | 330.0 | 1.0 |
| sam-lc-external | 0.02 | 0.57 | 0.0 | 0.0 |
| cp-lc-external | 0.15 | 0.44 | 80.0 | 0.24 |
| unet-sc-internal | 0.21 | 0.29 | 9.0 | 1.0 |
| d2-sc-internal | 0.32 | 0.37 | 8.0 | 0.89 |
| sam-sc-internal | 0.24 | 0.53 | 5.0 | 0.56 |
| cp-sc-internal | 0.11 | 0.21 | 3.0 | 0.33 |
| unet-lc-internallazy | 0.33 | 0.51 | 11.0 | 0.85 |
| d2-lc-internallazy | 0.41 | 0.52 | 12.0 | 0.92 |
| sam-lc-internallazy | 0.18 | 0.26 | 2.0 | 0.15 |
| cp-lc-internallazy | 0.44 | 0.49 | 5.0 | 0.38 |

### Comparing zero-shot and fine-tuning IoU performances

**Figure S2:**
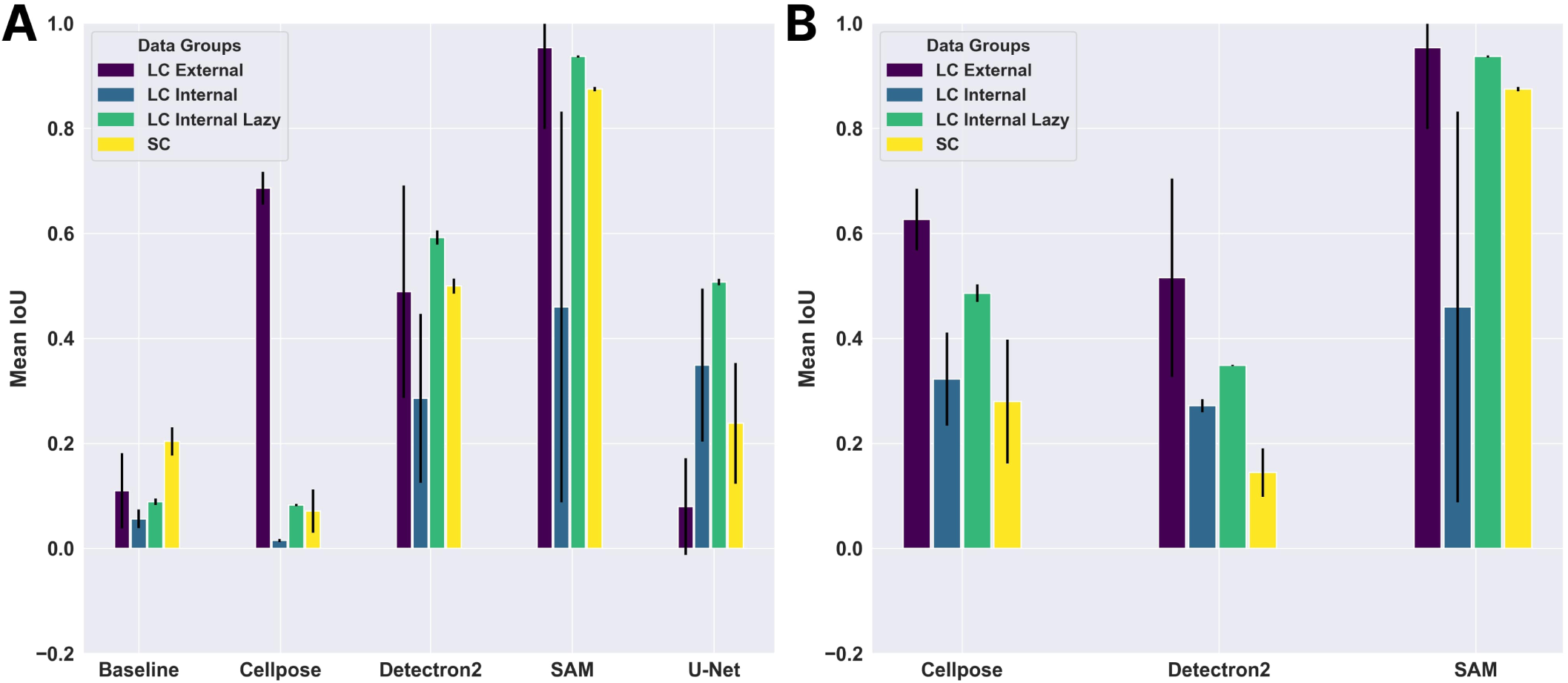
Figure A shows IoU for four fully-trained models and one baseline across all datasets for fine-tuning. Figure B shows the performance of zero-shot learning for models where zero-shot is applicable.

## REFERENCES

[1] Magdalena Strecanska, Tatiana Sekelova, Maria Csobonyeiova, Lubos Danisovic, and Michaela Cehakova. Therapeutic applications of mesenchymal/medicinal stem/signaling cells preconditioned with external factors: Are there more efficient approaches to utilize their regenerative potential? Life Sciences, 346:122647, 2024.

[2] Jacques Galipeau and Luc Sensébé. Mesenchymal stromal cells: Clinical challenges and therapeutic opportunities. Cell Stem Cell, 22(6):824–833, June 2018.

[3] Yusuke Shimizu, Edward Hosea Ntege, Chinatsu Azuma, Fuminari Uehara, Takashi Toma, Kotaro Higa, Hiroki Yabiku, Naoki Matsuura, Yoshikazu Inoue, and Hiroshi Sunami. Management of rheumatoid arthritis: Possibilities and challenges of mesenchymal stromal/stem cell-based therapies. Cells, 12(14):1905, July 2023.

[4] Global Health Datta. Gbd 2019: Global burden of 369 diseases and injuries in 204 countries and territories, 1990–2019: a systematic analysis for the global burden of disease study 2019. https://vizhub.healthdata.org/gbd-results/, 2019. Accessed: 2024-11-14.

[5] Maria Eugenia Fernández-Santos, Mariano Garcia-Arranz, Enrique J. Andreu, Ana Maria García-Hernández, Miriam López-Parra, Eva Villarón, Pilar Sepúlveda, Francisco Fernández-Avilés, Damian García-Olmo, Felipe Prosper, Fermin Sánchez-Guijo, Jose M. Moraleda, and Agustin G. Zapata. Optimization of mesenchymal stromal cell (msc) manufacturing processes for a better therapeutic outcome. Frontiers in Immunology, 13, June 2022.

[6] Dae Seong Kim, Myoung Woo Lee, Tae-Hee Lee, Ki Woong Sung, Hong Hoe Koo, and Keon Hee Yoo. Cell culture density affects the stemness gene expression of adipose tissue-derived mesenchymal stem cells. Biomedical Reports, 6(3):300–306, January 2017.

[7] Simon P. Shen, Hua-an Tseng, Kyle R. Hansen, Ruofan Wu, Howard J. Gritton, Jennie Si, and Xue Han. Automatic cell segmentation by adaptive thresholding (acsat) for large-scale calcium imaging datasets. eneuro, 5(5):ENEURO.0056–18.2018, September 2018.

[8] Costas Panagiotakis and Antonis A. Argyros. Cell segmentation via region-based ellipse fitting. In 2018 25th IEEE International Conference on Image Processing (ICIP), pages 2426–2430, 2018.

[9] Haoran Chen and Robert F. Murphy. Evaluation of cell segmentation methods without reference segmentations. Molecular Biology of the Cell, 34(6), May 2023.

[10] Olaf Ronneberger, Philipp Fischer, and Thomas Brox. U-Net: Convolutional Networks for Biomedical Image Segmentation, page 234–241. Springer International Publishing, 2015.

[11] Xu Han, Zhengyan Zhang, Ning Ding, Yuxian Gu, Xiao Liu, Yuqi Huo, Jiezhong Qiu, Yuan Yao, Ao Zhang, Liang Zhang, Wentao Han, Minlie Huang, Qin Jin, Yanyan Lan, Yang Liu, Zhiyuan Liu, Zhiwu Lu, Xipeng Qiu, Ruihua Song, Jie Tang, Ji-Rong Wen, Jinhui Yuan, Wayne Xin Zhao, and Jun Zhu. Pre-trained models: Past, present and future. AI Open, 2:225–250, 2021.

[12] Alexander Kirillov, Eric Mintun, Nikhila Ravi, Hanzi Mao, Chloe Rolland, Laura Gustafson, Tete Xiao, Spencer Whitehead, Alexander C. Berg, Wan-Yen Lo, Piotr Dollár, and Ross Girshick. Segment anything. In 2023 IEEE/CVF International Conference on Computer Vision (ICCV), pages 3992–4003, 2023.

[13] Yuxin Wu, Alexander Kirillov, Francisco Massa, Wan-Yen Lo, and Ross Girshick. Detectron2. https://github.com/facebookresearch/detectron2, 2019.

[14] Carsen Stringer, Tim Wang, Michalis Michaelos, and Marius Pachitariu. Cellpose: a generalist algorithm for cellular segmentation. Nature Methods, 18(1):100–106, December 2020.

[15] Christoffer Edlund, Timothy R. Jackson, Nabeel Khalid, Nicola Bevan, Timothy Dale, Andreas Dengel, Sheraz Ahmed, Johan Trygg, and Rickard Sjögren. Livecell—a large-scale dataset for label-free live cell segmentation. Nature Methods, 18(9):1038–1045, August 2021.

[16] Lucas von Chamier, Romain F. Laine, Johanna Jukkala, Christoph Spahn, Daniel Krentzel, Elias Nehme, Martina Lerche, Sara Hernández-Pérez, Pieta K. Mattila, Eleni Karinou, Séamus Holden, Ahmet Can Solak, Alexander Krull, Tim-Oliver Buchholz, Martin L. Jones, Loïc A. Royer, Christophe Leterrier, Yoav Shechtman, Florian Jug, Mike Heilemann, Guillaume Jacquemet, and Ricardo Henriques. Democratising deep learning for microscopy with zerocostdl4mic. Nature Communications, 12(1):2276, Apr 2021.

[17] R. Monarch, R. Munro, and C.D. Manning. Human-in-the-Loop Machine Learning: Active Learning and Annotation for Human-centered AI. Manning, 2021.

[18] Mohammad Jafari, Yimeng Zhang, Yihua Zhang, and Sijia Liu. The power of few: Accelerating and enhancing data reweighting with coreset selection. In ICASSP 2024 - 2024 IEEE International Conference on Acoustics, Speech and Signal Processing (ICASSP), page 7100–7104. IEEE, April 2024.

[19] Burcu Sayin, Evgeny Krivosheev, Jie Yang, Andrea Passerini, and Fabio Casati. A review and experimental analysis of active learning over crowdsourced data. Artificial Intelligence Review, 54(7):5283–5305, May 2021.

[20] Amit Kumar Gupta. Imglab.

[21] Tsung-Yi Lin, Michael Maire, Serge J. Belongie, Lubomir D. Bourdev, Ross B. Girshick, James Hays, Pietro Perona, Deva Ramanan, Piotr Doll’a r, and C. Lawrence Zitnick. Microsoft COCO: common objects in context. CoRR, abs/1405.0312, 2014.

[22] Tianzhixi Yin, Gihan Panapitiya, Elizabeth D. Coda, and Emily G. Saldanha. Evaluating uncertainty-based active learning for accelerating the generalization of molecular property prediction. Journal of Cheminformatics, 15(1), November 2023.

[23] Jingbo Zhu, Huizhen Wang, Tianshun Yao, and Benjamin K Tsou. Active learning with sampling by uncertainty and density for word sense disambiguation and text classification. In Donia Scott and Hans Uszkoreit, editors, Proceedings of the 22nd International Conference on Computational Linguistics (Coling 2008), pages 1137–1144, Manchester, UK, August 2008. Coling 2008 Organizing Committee.

[24] Henry B Mann and Donald R Whitney. On a test of whether one of two random variables is stochastically larger than the other. The annals of mathematical statistics, pages 50–60, 1947.

[25] John Canny. A computational approach to edge detection. *IEEE Transactions on Pattern Analysis and Machine Intelligence*, PAMI-8(6):679–698, 1986.

[26] Vishwesh Nath, Dong Yang, Bennett A. Landman, Daguang Xu, and Holger R. Roth. Diminishing uncertainty within the training pool: Active learning for medical image segmentation. IEEE Transactions on Medical Imaging, 40(10):2534–2547, October 2021.

[27] Daniel D Kim, Rajat S Chandra, Li Yang, Jing Wu, Xue Feng, Michael Atalay, Chetan Bettegowda, Craig Jones, Haris Sair, Wei-hua Liao, Chengzhang Zhu, Beiji Zou, Anahita Fathi Kazerooni, Ali Nabavizadeh, Zhicheng Jiao, Jian Peng, and Harrison X Bai. Active learning in brain tumor segmentation with uncertainty sampling and annotation redundancy restriction. Journal of Imaging Informatics in Medicine, 37(5):2099–2107, March 2024.

[28] Xiaokang Li, Menghua Xia, Jing Jiao, Shichong Zhou, Cai Chang, Yuanyuan Wang, and Yi Guo. Hal-ia: A hybrid active learning framework using interactive annotation for medical image segmentation. Medical Image Analysis, 88:102862, August 2023.

[29] Jiayu Huang, Nazbanoo Farpour, Bingjian J. Yang, Muralidhar Mupparapu, Fleming Lure, Jing Li, Hao Yan, and Frank C. Setzer. Uncertainty-based active learning by bayesian u-net for multi-label cone-beam ct segmentation. Journal of Endodontics, 50(2):220–228, February 2024.

[30] Wei Lou, Haofeng Li, Guanbin Li, Xiaoguang Han, and Xiang Wan. Which pixel to annotate: A label-efficient nuclei segmentation framework. IEEE Transactions on Medical Imaging, 42(4):947–958, 2023.

[31] Tianyi Zhao and Zhaozheng Yin. Weakly supervised cell segmentation by point annotation. IEEE Transactions on Medical Imaging, 40(10):2736–2747, October 2021.

[32] Rayan Krishnan, Pranav Rajpurkar, and Eric J. Topol. Self-supervised learning in medicine and healthcare. Nature Biomedical Engineering, 6(12):1346–1352, August 2022.

[33] Matthias Arzt, Joran Deschamps, Christopher Schmied, Tobias Pietzsch, Deborah Schmidt, Pavel Tomancak, Robert Haase, and Florian Jug. Labkit: Labeling and segmentation toolkit for big image data. Frontiers in Computer Science, 4, February 2022.

[34] Nikhila Ravi, Valentin Gabeur, Yuan-Ting Hu, Ronghang Hu, Chaitanya Ryali, Tengyu Ma, Haitham Khedr, Roman Rädle, Chloe Rolland, Laura Gustafson, et al. Sam 2: Segment anything in images and videos. *arXiv preprint arXiv:2408.00714*, 2024.

[35] Sushish Baral and May Phu Paing. Instance segmentation of cells and nuclei from multi-organ cross-protocol microscopic images. Quantitative Imaging in Medicine and Surgery, 14(9):6204–6221, September 2024.

[36] Rajan Gyawali, Ashwin Dhakal, Liguo Wang, and Jianlin Cheng. Cryosegnet: accurate cryo-em protein particle picking by integrating the foundational ai image segmentation model and attention-gated u-net. Briefings in Bioinformatics, 25(4), May 2024.

[37] Anwai Archit, Sushmita Nair, Nabeel Khalid, Paul Hilt, Vikas Rajashekar, Marei Freitag, Sagnik Gupta, Andreas Dengel, Sheraz Ahmed, and Constantin Pape. Segment anything for microscopy. August 2023.

